# Constraint Induced Movement Therapy Confers only a Transient Behavioral Benefit but Enduring Functional Circuit-Level Changes after Experimental TBI

**DOI:** 10.1101/2024.08.02.606449

**Authors:** Afshin Paydar, Laila Khorasani, Neil G. Harris

**Affiliations:** UCLA Brain Injury Research Center, Department of Neurosurgery, Geffen Medical School; Intellectual Development and Disabilities Research Center, University of California at Los Angeles, Los Angeles, CA, 90095, USA

**Keywords:** Traumatic Brain Injury, Reorganization, Constraint-Induced Movement Therapy, rehabilitation, resting state fMRI, Functional Connectivity, fMRI

## Abstract

Although the behavioral outcome of Constraint-Induced Movement Therapy (CIMT) is well known, and that a combination of CIMT and arm use training potentiates the effect, there has been limited study of the brain circuits involved that respond to therapy. An understanding of CIMT from a brain network level would be useful for guiding the duration of effective therapy, the type of training regime to potentiate the outcome, as well as brain regional targets that might be amenable for direct neuromodulation. Here we investigated the effect of CIMT therapy alone unconfounded by additional rehabilitation training in order to determine the impact of intervention at the circuit level. Adult rats were injured by controlled cortical impact injury and studied before and then after 2wks of CIMT or noCIMT at 1-3wks post-injury using a combination of forelimb behavioral tasks and task-based and resting state functional magnetic resonance imaging at 3 and 7wks post-injury and compared to sham rats. There was no difference in behavior or functional imaging between CIMT and noCIMT after injury before intervention so that data are unlikely to be confounded by differences in injury severity. CIMT produced only a transient reduction in limb deficits compared to noCIMT immediately after the intervention, but no difference thereafter. However, CIMT resulted in a persistent reduction in contralesional limb-evoked activation and a corresponding ipsilesional cortical plasticity compared to noCIMT that endured 4wks after intervention. This was associated with a significant amelioration of intra and inter-hemispheric connectivity present in the noCIMT group at 7wks post-injury.

## Introduction

Motor weakness occurs in 9-56% of patients following traumatic brain injury (TBI)(Katz, Alexander, and Klein 1998; Gagnon et al. 1998; Tremblay et al. 2011; Hillier, Sharpe, and Metzer 1997; De Beaumont et al. 2009) with up to 30% showing persistent deficits in upper limb function 2-5 years post-TBI (Hillier, Sharpe, and Metzer 1997). Deficits in sensory-motor cortical inhibition have been shown even in mild TBI over 1 year following a single or multiple concussions (Brown et al. 2007). Despite good spontaneous recovery in patients following injury, abnormal movement-related cortical potentials often remain (Di Russo et al. 2005). Persistent motor deficits are an important sequela of TBI, but as yet they have received little attention as a target of clinical interventional strategies and experimental, mechanistic research.

Prior preclinical research has shown that the contralesional cortex opposite the primary injured hemisphere is causally involved in deficient limb function (Verley et al. 2019). At least over the first week after injury in adult rats, silencing the contralesional cortex rescues limb function; however, by 4 weeks, the same neuromodulation worsens limb use (Verley et al. 2019). The prevailing theory for this is that hemispheric imbalance in excitation-inhibition as supported by temporally increasing excitability of the contralesional cortex after injury (Verley et al. 2018), prevents functional connectivity to the ipsilesional forelimb, pericontusional cortex. The observed early behavioral improvement is underpinned by neuromodulatory-induced increases in functional connectivity across the brain at the cortical level, and between the pericontused cortex bilaterally to the thalamus. The fact that these circuit changes are absent at 4wks post-injury coincident with failure to enhance limb function (Verley et al. 2019), indicates a shift in circuit plasticity overtime that prevents acute neuromodulatory control. While the pericontused cortex temporarily remains active to input after injury, the route of activation indicates that the ipsilesional pericontused cortex is functionally isolated from normal afferent input (Paydar and Harris 2023). The question remains whether a more temporally extended modulation of function would overcome this block in functional activation from the opposite, affected forelimb and promote more permanent limb recovery function. In this current work, we have used 2weeks of constraint-induced movement therapy (CIMT) to test whether prolonged intervention to force use the injury- affected limb would alleviate the limb use deficit.

The CIMT rehabilitation technique has been mostly applied and studied in stroke rather than TBI patients, although there are a number of exceptions (Greenwald and Rigg 2009; Pedlow, Lennon, and Wilson 2014; Shaw et al. 2005; Zhao et al. 2012; Cimolin et al. 2012). CIMT aims to correct the maladaptive preference to use the less-affected limb that suppresses the use of injury-affected limb by limiting the less-affected limb, forcing use of the injury-affected limb. Current evidence suggests that CIMT combined with forced repetitive use of the injury–affected forelimb can have beneficial effects on limb function after TBI (Greenwald and Rigg 2009; Pedlow, Lennon, and Wilson 2014; Shaw et al. 2005; Zhao et al. 2012; Cimolin et al. 2012). CIMT has been studied preclinically in rats after moderate to severe TBI where it, when paired with rehabilitative training or excise, shows a significant improvement in limb function. (Adkins et al. 2015; Combs et al. 2016; Lam et al. 2013). Here, we sought to determine whether a prolonged period of CIMT alone and uncomplicated by other rehabilitative effects would be enough to promote an alteration in neural circuits relating to forelimb function, and whether this would be accompanied by some measurable, contemporaneous enhancement in limb function. We postulated that, given that the forelimb pericontused cortex remains amendable to forelimb activation early after injury (Paydar and Harris 2023), any improvement in injury- affected limb function would involve a reconfiguration of functional circuits relating to the ipsilesional hemisphere, in much the same way that a direct neuromodulation of the cortex would (Paydar and Harris 2023).

In this study, we used a mild/moderate controlled cortical impact in adult rats and studied the effects of CIMT on forelimb functional circuitry using forelimb behavioral task outcomes together with resting state and fore-limb evoked functional MRI both after injury before intervention, or injury and sham-injury with no intervention, as well as after CIMT intervention. While a temporary improvement in limb use was conferred by 2wks of CIMT intervention in agreement with prior work, the major findings relate to enduring functional evoked and connectivity changes. Contralesional cortical map changes occurred as a direct result of lack of use of the less-affected limb, indicating a brain-centric effect of limb constraint which endured even after intervention. Ipsilesional map plasticity persisted through forced use of the injury-affected limb, while injury-associated loss of some transhemispheric cortico-cortical and corticothalamic connectivity was prevented by CIMT intervention and included parts of the pericontused cortex. These data comprise a novel and fundamental understanding of the effects of CIMT at a circuit level, and could provide a framework with which to understand how combinatorial rehabilitation paradigms can optimize these circuit changes to improve outcome.

## Materials and Methods

### Experimental Groups and Procedure

Adult, male, Sprague-Dawley rats at 250– 300g either received a controlled cortical impact (CCI) brain injury (n=30) or were used as anesthesia + scalp incision, sham controls (n=15). CCI rats were randomly assigned to injured+CIMT or injured+noCIMT. CIMT was administered from 1- 3wks after injury. Behavioral and functional magnetic resonance imaging (fMRI) data were acquired longitudinally before, during and after CIMT (**Fig. 1**). Rats were housed in pairs before and after the surgery in standard cages with *ad libitum* access to water and food. Rats were kept in regular 12:12 hour light/dark cycle in vivarium. All study protocols used were in compliance with the Public Health Service Policy on Humane Care and Use of Laboratory Animals and approved by the Chancellor’s Animal Research Committee, University of California, Los Angeles (UCLA) Approval #2021-082.

**Fig. 1.**
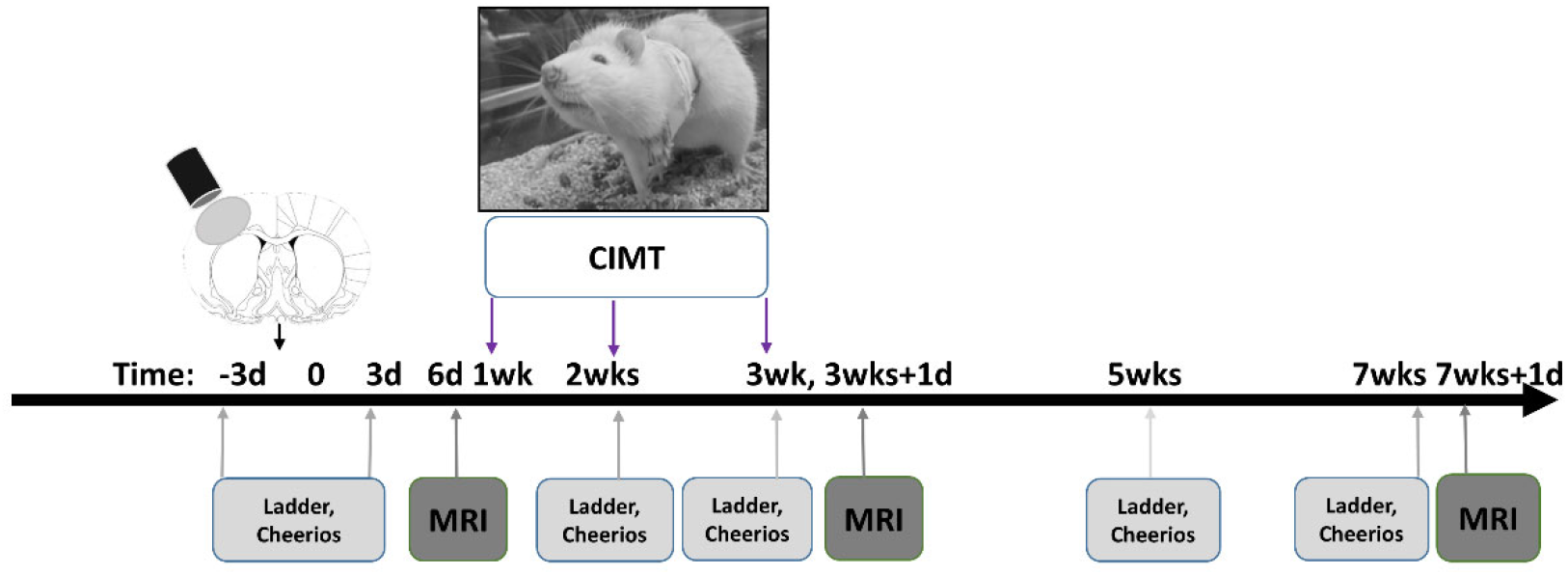
Study Timeline. Adult, Sprague Dawley rats received a CCI injury over the left somatosensory cortex. Following a post-injury behavioral assessment using the ladder walk and cheerios task and 1 session of functional MRI, constraint-induced movement Therapy (CIMT) was applied continuously for 2 weeks to the left forelimb (CIMT-targeted limb) to force-use the right, injury-affected limb. Two more behavioral tests were conducted at 2 and 3wks postinjury during CIMT and at 5 and 7wks post-injury once CIMT was completed. MRI data were acquired immediately following CIMT and at 7wks post-injury.

### Brain Injury Surgery

Moderate CCI injury was induced based on previously described methods^19–21^. In brief, rats were anesthetized with 5% isoflurane vaporized in 1 L/min O2 (2% isoflurane was used for maintenance of the anesthesia), and then were head-fixed in a stereotactic frame and their body temperature was maintained at 37°c using a temperature-controlled heating-pad. A 5mm craniotomy (centered at AP: +0.5mm and 3.5mm left side of the Bregma) was made and CCI injury was done by a 4mm diameter flat metal impactor tip onto the brain using a pneumatic pressure system at 20-psi and an impact depth of 2 mm lower than the dura surface with a dwelling time of 250 ms. The craniotomy defect was covered with a layer of biologically inert elastomer KWIK-Cast silicone sealant (World Precision Instruments, Sarasota, FL, USA).

### Ladder Rung Walking Task

To evaluate sensorimotor coordination rats were tested for their ability to walk on a horizontal metal ladder with a 2 cm distance between rungs. Each session was video recorded and then manually scored to calculate the affected, right forelimb fault score using this formula: Fault score=full forelimb faults + 0.5*half forelimb Faults/total steps*100.

This test was repeated at several time-points: before the injury (pre-injury), 3 days after the CCI injury (PID 3), after a week of the CIMT intervention (PID 14), after 2 weeks of the CIMT treatment (PID 21) and 4 weeks after the end of the CIMT intervention (PID 49) (Fig. 1).

### Cheerio Task

To assess fine forelimb sensorimotor skill we use the cheerio task with some modifications based upon similar tests in the literature (Irvine et al. 2010; Walker et al. 2012). Rats were first habituated to the task over 7 days by providing them with 5-6 cheerios in addition to their daily chow. For the test each rat was placed inside a transparent plexiglass hollow cylinder and provided with 10 cheerios (Honey Nut Cheerios) for consumption. Their behavior was recorded using a video recorder that allowed the subject to be viewed at high magnification for scoring after the task was completed. We developed a scoring system based on previous studies to evaluate their ability to hold and maneuver each cheerio while eating, based on parameters such as: keeping the cheerio off the ground, grasping the cheerio with palmar and digit support, and successful manipulation of the cheerio. Scores were tallied based on a point system with a maximum of 50 points (**Table 1**).

**Table 1.** Cheerios Task Scoring System (a higher total score indicates fewer deficits in fine sensorimotor skills).

|  | **Drops** | **Maintain**<br>**forelimbs off**<br>**ground &**<br>**limbs**<br>**evenness** | **Treat**<br>**manipulation**<br>**(affected limb)** | **Volar support**<br>**(affected**<br>**limb)** | **Digit contact**<br>**(affected**<br>**limb)** |
| --- | --- | --- | --- | --- | --- |
|  | No Drop=1 | Even + above<br>ground =1 | Good<br>adjustment +<br>full control =1 | Palm contact<br>=1 | Full =1 |
|  | Drop=0 | Uneven +<br>above ground<br>=0.5 | Rely on<br>unaffected limb<br>=0.5 | Palm contact<br>=0 | Partial =0.5 |
|  |  | On ground<br>completely =0 | No adjustment<br>+0 |  | None =0 |
| **Cheerios**<br>**count** | x10 | x10 | x10 | x10 | x10 |
| **Maximum** |  | **50** |  |  |  |
| **Total score** |  |  |  |  |  |

### Constraint-Induced Movement Therapy (CIMT)

On day 7 post-injury, rats were placed in rat jackets (Lomir Inc., Malone, NY, USA) and their ipsilesional (left or less-affected forelimb) was covered with athletic bandage such that the rats couldn’t use their less-affected forelimb for walking around their cage or eating, and would be required to use their injury-affected (right) forelimb. CIMT was applied continuously for 2 weeks from day 7 to the end of day 20 post-injury (**Fig. 1**) and rats were inspected several times every day to make sure that the jacket and the bandage remained in place.

### MRI Image Acquisition

MRI data were acquired at 6 days (before starting CIMT intervention), 21 days (after the end of CIMT), and 49 days after the injury. A 7 Tesla Bruker magnet with a Bruker ‘S116’ gradient insert (400mT/m), a transmit coil, and a receiver surface coil were used for data acquisition. During the MRI scanning, rats were sedated using dexmedetomidine hydrochloride (Dexdomitor, Orion Pharma, Espoo, Finland; Zoetis Inc. Kalamazoo, MI, USA with an intravenous bolus of 0.05mg/kg, and then a continuous subcutaneous infusion of 0.1mg/kg/hr). Their body temperature was kept constant at 37±- 0.5°C using a hot air heater controlled by an animal body temperature feedback system, and their respiratory rate was monitored (SA Instruments, Stony Brook, NY). After the set- up, an anatomical T2-weighted Rapid-Relaxation-with-Enhancement (RARE) image (50x0.5mm slices, 128x128 in a 35x35mm field-of-view); TR /TE=5000/60, RARE factor 8 was obtained. Resting state FMRI (rsFMRI) data were acquired for 15 minutes (450 volumes) using a single-shot, echo-planar, gradient echo sequence with a 128 x 64, read x phase data matrix in a 30x30mm field of view, and with 14 x 0.75mm slices, repetition/echo time (TR/TE) =2000/30ms), 90° flip angle. The same gradient echo sequence was used to acquire forelimb-evoked, task-based fMRI data, before, during and after 10 periods of forelimb electrical stimulation (30s ‘OFF’-20s ‘ON’) for each forelimb separately. Each forelimb was electrically stimulated through a pair of subcutaneously-placed needle electrodes using a –2mA, 10Hz stimulation, a level below the threshold for activation of pain fibers, similar to our previous studies. At the end of each imaging session, the rats were recovered using the dexmedetomidine antagonist, atipamezole hydrochloride (Antisedan, Orion Pharma, Espoo, Finland; Zoetis Inc. Kalamazoo, MI, USA; 1.0mg/kg, intraperitoneally).

### MRI Data Analysis & Statistics

We used an in-house, python parallel data processing pipeline for rsfMRI data pre-processing using command line code from the FMRIB Software Library (Smith et al. 2004), Advanced Normalization Tools (ANTS) (Avants et al. 2014), and the Rapid Automatic Tissue Segmentation (RATS) for rodent brain extraction (Oguz et al. 2014). Briefly, rsFMRI data preprocessing include these steps: slice timing correction; motion correction; brain extraction; segmentation into grey matter, white matter and CSF to exclude all voxels containing CSF; movement regression of 6 movement parameters from the data; 0.8mm Gaussian smoothing; band-pass filtering between 0.01-0.25Hz; registration to the corresponding brain extracted anatomical data and subsequently to a parcellated rodent brain atlas used previously (Harris et al. 2016). Time-course data were extracted from 30 regions encompassing sensory and motor cortices and subcortical regions of thalamus and striatum and were used to derive functional connectomic matrices between each region using Pearson correlations. After converting correlation values to z scores via Fisher transformation, regional edgewise differences between groups were tested for using a 2-tailed general linear model design with permutation testing (5000 shuffles), and multiple comparison correction using network-based statistics (P<0.05) implemented using the BrainNetwork Toolbox (Rubinov & Sporns, 2010) under GraphVar (Kruschwitz et al., 2015).

Task-based fMRI data were post-processed in the same pipeline for slice timing correction; motion correction; brain extraction before fitting the to the square wave of the repeating OFF/ON 30s/20s stimulation sequence convolved with a Gamma waveform of phase=0s, standard deviation=0.5s and a mean lag time of 3s, and Gaussian smoothed to 0.2mm. Mixed effects modelling using cluster-based thresholding correction at z=1.7, P<0.05 was used to assess the group differences for the data (Worsley 2001).

To evaluate the local tissue deformations for approximating tissue atrophy/compression and swelling/expansion, we used anatomical data similar to previous studies^25^. In brief, a mean deformation brain template was constructed from all data using ANTS^31^ and then the Jacobian Determinants from the resulting warp transformation fields were calculated. After calculating voxel-based mean and standard deviation maps from the sham group Jacobian data which used to calculate z statistic maps for all injured groups data. For each injured brain, the volume of tissue surviving a threshold of z ±3.01 (P<0.01) was determined and then used to assess the region of local tissue expansion and contraction.

Behavioral measures were tested for significance using a longitudinal design and a mixed model analysis of variance, with posthoc testing adjustment for multiple comparison implemented using Tukey’s test (P<.05) under Graphpad, Prism (v8). We tested for normality of the data distribution using the D’Agostino-Pearson test and since data at 5 and 7 wks post-injury data did not pass, all data were log-transformed before further analysis. We explicitly split the analysis into two longitudinal segments: (i) pre-injury to post-injury before-CIMT and (ii) post- CIMT in order to test for the effect of injury alone, and the effect of CIMT, respectively.

## Results

To simplify the nomenclature when describing the left versus right forelimb in relation to left sensorimotor cortex injury and left forelimb CIMT, we refer to the left cortical hemisphere, the primarily injured hemisphere, as the ipsilesional hemisphere, and the right hemisphere, opposite to the primarily impacted hemisphere as the contralesional hemisphere. In addition, we refer to the left forelimb which is controlled by the right (contralesional) cortex and is the target for CIMT as the “less-affected forelimb”. The right forelimb that is predominantly affected by the left side unilateral injury is referred to as the “injury-affected forelimb”. (**Fig. 1**).

### No significant difference among injured groups before CIMT indicating matched injury severities

We explicitly analyzed the data in two separate longitudinal sections: before and after CIMT, in order to first test for the existence of any differences between the injury groups prior to intervention that might confound the interpretation of any potential therapeutic effect. The effect of injury on forelimb behavioral function was similar in both injured-CIMT and noCIMT groups, where injury resulted in significant alterations in both gross and fine motor skills compared to sham before CIMT at 3d post-injury (P<.001), but no difference between the injured groups (**Fig.2A,B**). Direct testing of affected forelimb function using fore-limb-evoked BOLD activity showed similar effects to behavior, with significant decreases in cortical activity in both injured groups compared to sham before CIMT at 1wk post-injury, and only minor differences between injured groups in regions outside of the forelimb circuit (**Fig. 2C**). There was no injury group difference in cortical activity evoked by stimulation of the less-affected limb (**Fig. 2D**). Resting state functional connectivity (fc) data acquired at the same 1wk post-injury imaging session showed similar effects, with mainly transhemispheric deficits in fc in both injured groups compared to sham, with very minor differences between the injured groups (**Fig. 2E-G**). These fc data reflect largely thalamocortical circuity likely to be involved in both sensory cortex injury. Whole brain tensor-based deformation analysis of the T2-weighted, anatomical data at 7wks post-injury revealed similar spatial z-scores across the brain in both injured groups compared to sham (**Fig. 2H,I**). A quantitative volume analyses of these data at 1, 3 and 7wks showed that there were no differences in either local tissue expansion or compression between the injured groups (p>.05) either before or after CIMT (**Fig. 2J,K**). The combination of these behavioral and brain functional and structural data therefore indicate that the injury levels were comparable across the injured groups, so that any differences in behavior, brain activation and circuit connectivity are unlikely to be confounded by differences in injury severity.

**Fig. 2.**
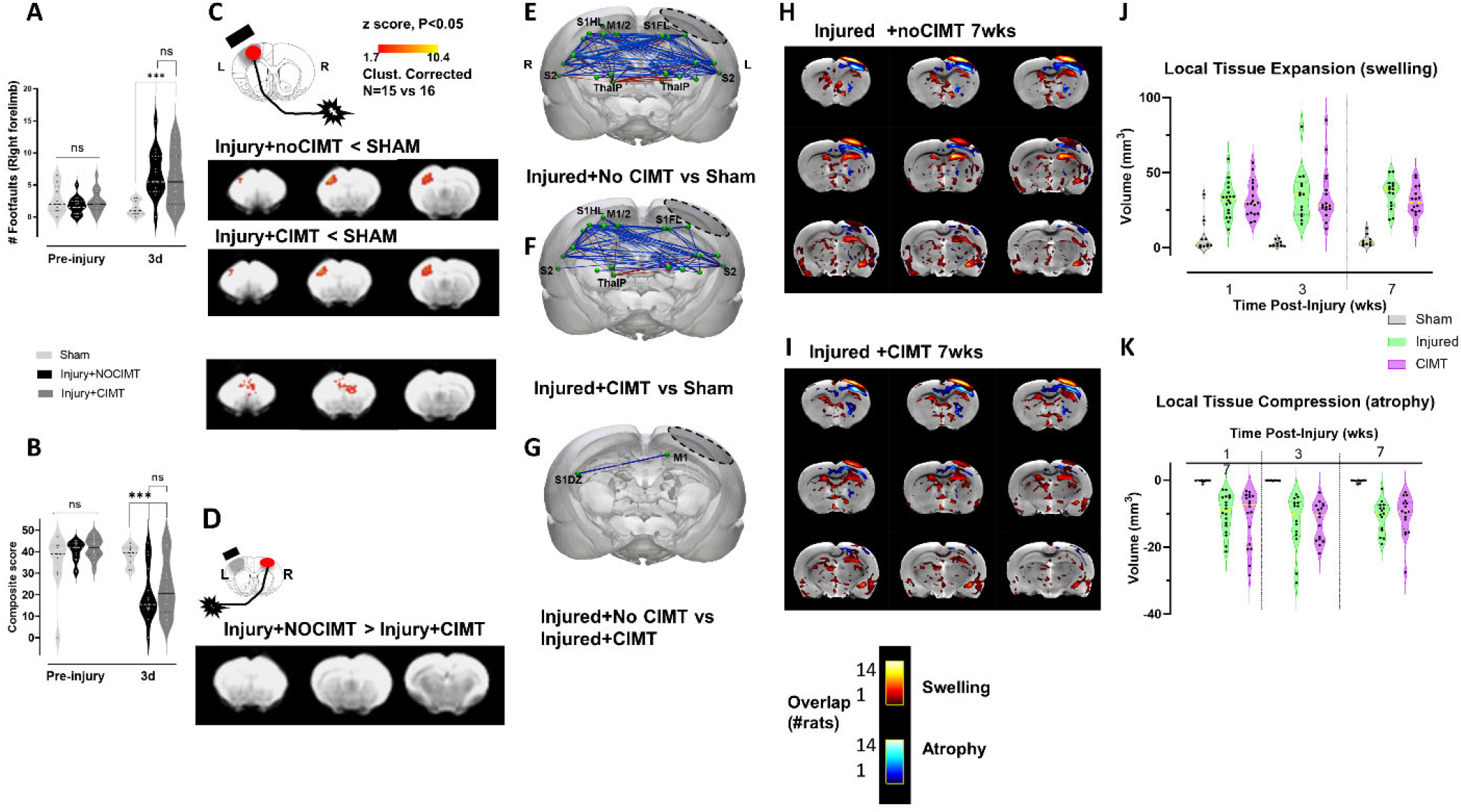
Absence of differences in behavior [A-B], evoked activity [C-D] and functional connectivity [E-G] between injured groups before beginning CIMT and structural changes chronically[H-k] indicating matched injury severity. [**A**] The ladder task of gross sensorimotor skills showed a significant increase in right, injury-affected limb foot faults at 3d after injury+CIMT and NOCIMT compared to shams but there was no difference in performance among the injured groups prior to beginning CIMT. [**B**] The cheerios task of fine sensorimotor skills showed a significant decrease in both injured groups compared to preinjury at 3d post-injury, but no difference between the groups before beginning CIMT. [**C**] While cortical evoked activation was reduced in both injured groups compared to sham when elicited by limb stimulation opposite the primary injury at 1wk post-injury, a direct comparison between injured groups revealed minor differences in prefrontal and cingulate region activation, but not in sensory motor regions. [**D**] There were no evoked activity differences between injured groups when the less-affected, left limb ipsilateral to the injury was stimulated. [**E-F**] Functional connectivity by resting state FMRI at 1wk after injury and before CIMT, resulted in transhemispheric hypoconnectivity between the two cortices (blue edges) and hyperconnectivity between the left ipsilesional cortex and the contralesional subcortical regions (red edges) in both Injured-no CIMT and injured+CIMT versus the sham group (P<.05, Network-based statistics) [**G**] However, there were only minor differences between the injured group on direct comparison. [**H-I**] Z-map incidence plots from each injured rat thresholded at P < 0.01 and binarized to show the overlap in regions of local tissue expansion/swelling (red/yellow) and compression/atrophy (blue) compared to sham at 7wks after injury in injured+CIMT and injured-no CIMT groups. [**J-K**] There were no significant group differences in the volume of local tissue compression and/or expansion between the two injured groups (p>0.05) assessed by Jacobian z-score data, indicating that differences in brain activation and circuit connectivity are unlikely to be confounded by differences in injury severity. Key: ***=P<.001, ns=not significant. L=left, R=right. <u>Brain Regions</u> (E-F): S1HL- sensory hindlimb cortex, S1FL- sensory forelimb cortex, M1/M2- motor cortical regions, S2- secondary somatosensory cortex, ThalP- posterior thalamus, S1DZ- sensory dysgranular cortex (See Fig. 5 for additional labels). Left Dotted Boundary Region- primary impact area.

### CIMT only transiently improved gross motor skills but the effect on the cortical-evoked map was persistent

After injury, CIMT was conducted 24hrs/day for 2wks from 1-3wks post-injury and we tested for improvement in gross and fine motor skills. The effect of CIMT was only transient and trending toward significantly fewer foot faults measured immediately after the CIMT intervention when compared to injured+noCIMT (P=0.059, corrected). The absence of significance was driven by a single rat that was unresponsive to CIMT. However, the effect of injury alone had significantly diminished by this time of 3wks after injury so that there was no longer a difference compared to sham (P>.05), **Fig. 3A**). Thereafter the effect of CIMT was not different from noCIMT, and the effect of injury also did not endure relative to sham. Curiously, there was a worsening of gross motor skills at 5wks due to CIMT but not in noCIMT compared to sham (P<.05), but this was only transient. The effect of injury on fine motor skills resolved by 2wks post-injury so that there were no significant effects compared to sham or due to CIMT (**Fig. 3B**). Despite these transient and lackluster behavioral effects, the cortical map evoked by stimulation of the constrained/less-affected limb immediately after the 2wk CIMT period was statistically reduced compared to the injured+noCIMT group (P<.05, **Fig.4A**), clearly indicating a brain-centric effect of limb constraint on map plasticity. Moreover, this effect persisted 4wks later, after the end of the CIMT period at 7wks post-injury (**Fig. 4A**).

**Fig. 3.**
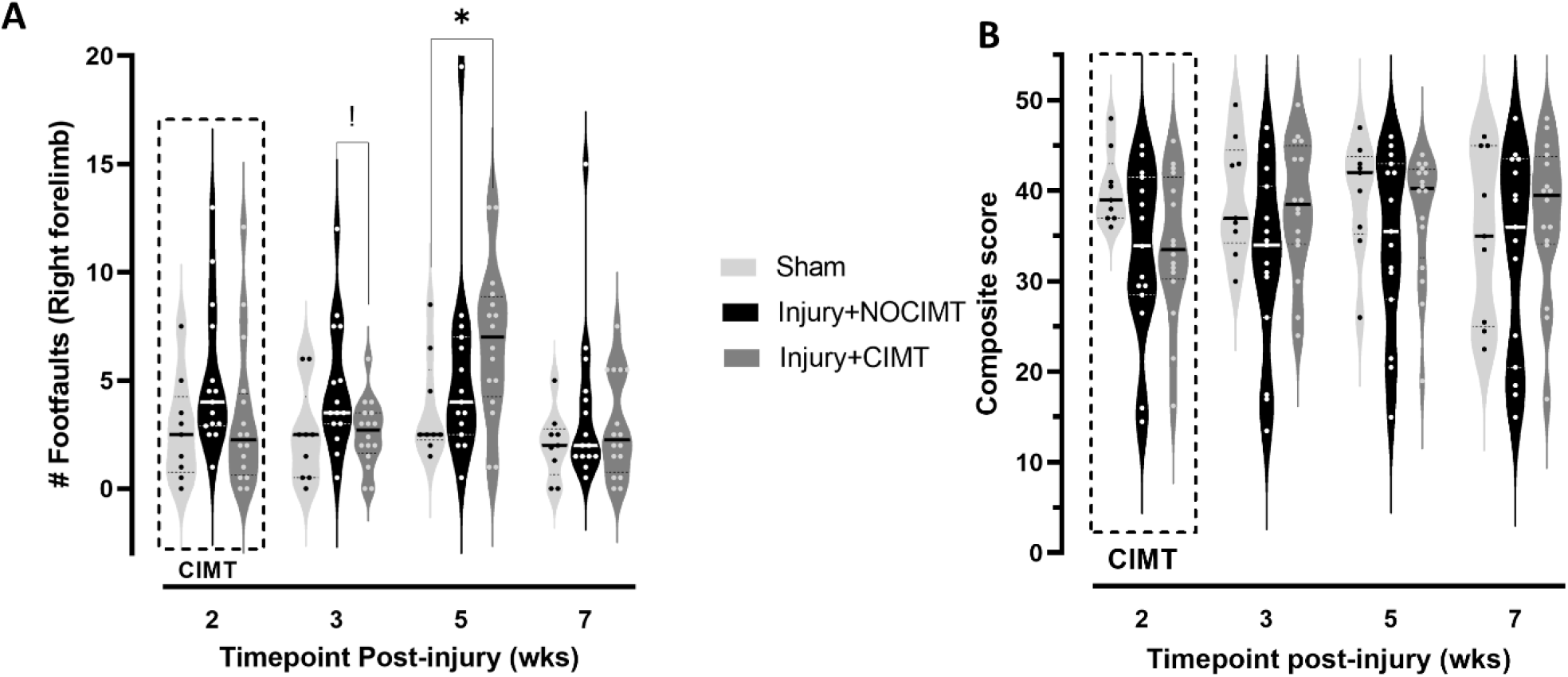
Behavioral Effect of CIMT Intervention on gross and fine sensorimotor skills with the [A] ladder task of gross sensorimotor skills and [B] The cheerios task of fine sensorimotor skills. CIMT intervention at 1-3wks post-injury resulted in [**A**] a temporary but significant reduction in foot faults on the ladder task at 3wks immediately after CIMT compared to injured+noCIMT. However, by 5wks the beneficial effect of CIMT was lost, and it temporarily resulted in increased numbers of faults compared to sham. [**B**] The effect of injury and injury+CIMT resulted in no discernable long-term deficits in fine motor skills. Two-way mixed effects ANOVA with Tukey’s correction for multiple comparisons. Key: *=P<.05, != P=.059

**Fig. 4.**
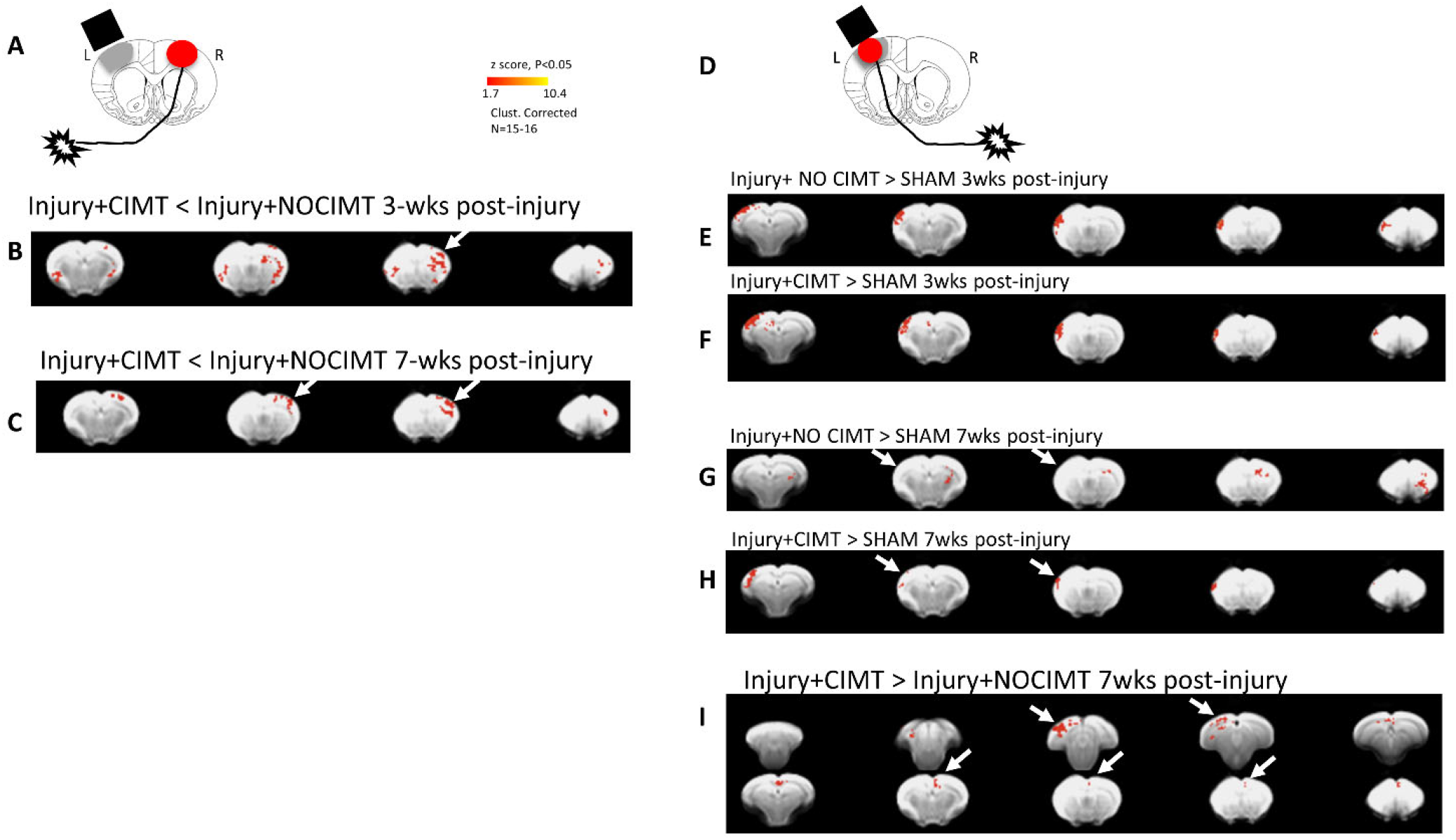
CIMT-induced Cortical-Evoked Map Plasticity. [**A**] CIMT of the less-affected limb [**B**] persistently reduced forelimb-evoked responsivity of the contralesional cortex at 3wks and [**C**] this endured even 4wks after completion of CIMT at 7wks post-injury (arrows). [**D**] The effect of CIMT intervention on forced use of the affected limb contralateral to the injury was first measured against sham. The difference of each injury group to sham at [**E-F**] 3wks and [**G-H**] 7wks indicated a persistent ipsilateral S2 map plasticity at 7wks after injury+CIMT but not in the injured+no-CIMT group where it was present only at 3wks (arrows). [**I**] Direct comparison of the two injury groups revealed subtle but persistent increases in map plasticity at 7wks post-injury, 4wks following the end of CIMT intervention (arrows).

### CIMT Rehabilitation induces persistent remapping of Ipsilesional Cortical activity

The cortical evoked map elicited by stimulation of the affected limb opposite the primary injury was similarly increased from shams in both injured groups at 3wks, immediately after CIMT in both cortical S2 and posterior cortical regions outside the normal forelimb active region (**Fig. 4D,E,F**). There was no difference between the injured groups due to CIMT at this time (not shown). However, while this map plasticity waned with time in injured+noCIMT rats at 7wks post-injury (**Fig. 4G**), the effect persisted in injured+CIMT rats (**Fig. 4H**). Direct comparison of the two injured groups at 7wks demonstrated this enduring difference was localized to both anterior cingulate and M2 regions, but also to posterior, ipsilesional cortical regions (**Fig. 4I**).

### Functional connectivity (FC) changes reflect CIMT Intervention

Injury alone resulted in mainly persistent transhemispheric disconnection relative to shams within the forelimb circuit, together with the development of some thalamocortical hyperconnectivity at both 3 and 7wks (**Fig. 5**). Injury+CIMT was associated with a significant attenuation of these FC patterns compared to sham. The initial direct effect of CIMT on FC was characterized by strengthened transhemispheric motor and sensory S2 communication, and a reduction in sensory S1 and thalamic connectivity at 3wks (**Fig. 5**). This effect was replaced 4 weeks later by persistent increased FC emanating solely between ipsilateral sensory hindlimb cortex and bilateral sensory and motor regions, as well as contralateral thalamus.

**Fig. 5.**
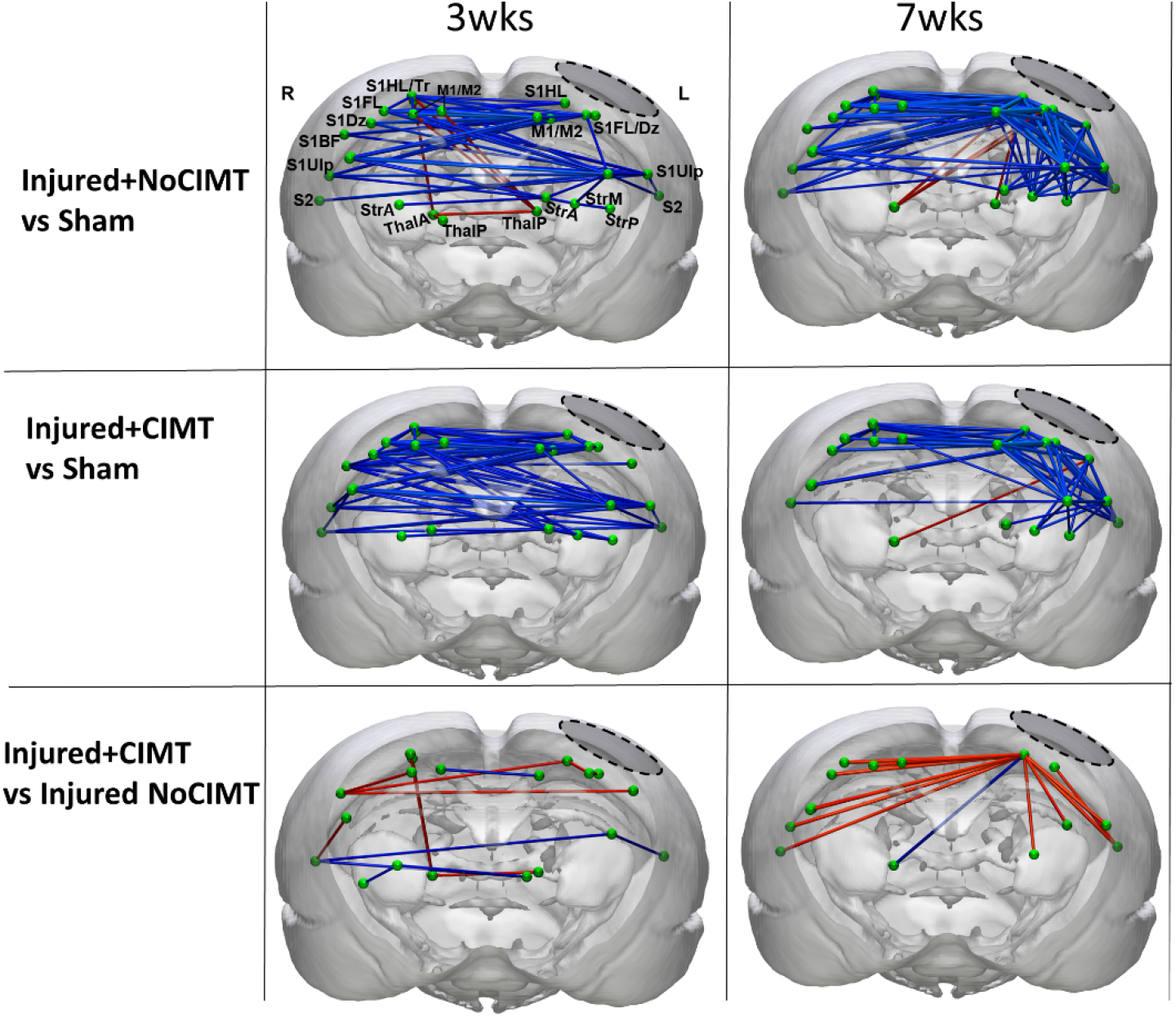
CIMT-induced Alterations in Functional connectivity. [**Upper and Middle Rows**] Injury, regardless of CIMT intervention, resulted in transhemispheric hypoconnectivity between the two cortices (blue edges) and hyperconnectivity between the left ipsilesional cortex and the contralesional subcortical regions (red edges) of Injured-no CIMT and injured+CIMT versus the sham group at 3 and 7wks post-injury. [**Lower Row**] Direct comparison of the two injured groups revealed CIMT-induced increases and decreases among mainly cortical sensory and motor and thalamic regions at 3wks after injury directly at the end of the CIMT period, and transhemispheric increases in mainly cortical sensory regions and ipsilateral striatum at 7wks that persisted 4wks after the end of the CIMT intervention. Data were computed using a 2-tailed general linear model using 5000 permutation with network based statistical correction (P<0.05). Blue and red edges indicate decreases increases, respectively. <u>Brain Regions</u> (E-F): S1HL- sensory hindlimb cortex, S1FL- sensory forelimb cortex, M1/M2- motor cortical regions, S2- secondary somatosensory cortex, ThalP- posterior thalamus, S1DZ- sensory dysgranular cortex, S1TR- sensory trunk cortex, S1BF- barrel cortex, S1UP- sensory upper lip, StrA/M/P- Anterior, Medial or Posterior Striatum, ThalA/P- Anterior or Posterior Thalamus. Left Dotted Boundary Region- primary impact area.

## Discussion

The major goal of this study was to determine whether there are specific circuit-related changes that accompany the effect of limb constraint therapy after injury. By identifying these potential effects, we might then begin to understand how future work combining the effect of other rehabilitation paradigms, such as those used previously with CIMT in rodents (Adkins et al. 2015; Combs et al. 2016) could be used in a focused way to target particular circuits beneficial to outcome. Importantly, we were able to discern the direct functional effects of CIMT on the cortex opposite the constrained limb using a cortical evoked fMRI paradigm, and we found the effect was longer lasting than simply the 2-week period of limb constraint. That is, the lack of use of the less-affected limb produced an enduring loss of evoked activity in the opposite cortex, contralesional to the primary injury site. The indirect effect of this was to enhance ipsilesional cortical map plasticity evoked by the stimulation of the opposite, injury-affected limb, and once again this effect endured 4weeks past the end of the CIMT intervention. These functional changes were driven by only very mild, non-significant improvements in forelimb use, as determined by behavioral measures that were only temporarily trending different from unconstrained, injured rats. Despite this, the 2week CIMT intervention did produce a robust improvement in transhemispheric functional connectivity as well as a reduction in corticothalamic hyperconnectivity.

The lack of robust behavioral effects with CIMT alone found in this study is broadly in agreement with prior work in rats in which only a combination of interventions resulted in improved limb use (Adkins et al. 2015; Combs et al. 2016). The novel findings in this work relate to the enduring pliability of the forelimb functional circuits due to limb constraint that show similarities to direct cortical neuromodulation (Paydar and Harris 2023). Acute silencing the contralesional cortex with the GABA agonist muscimol at 1wk after injury in the same TBI model reinstates injury-affected limb use and drives significant enhancement in cross-brain fc between both sensory and motor regions in much the same way as CIMT does in the current work. However, there the similarities end since acute direct contralesional neuromodulation did not reinstate forelimb-evoked activity of the ipsilesional cortex even when administered at 1wk after injury (Paydar and Harris 2023), while CIMT does result in ipsilesional sensory S2 region BOLD activity that endures for many weeks after the intervention as well as scattered posterior cortical activation. While these two datasets are based upon very different therapeutic windows for neuromodulation: ∼2hrs muscimol versus 2wks of CIMT, the similarities in FC changes do indicate similar circuits are modulated, while the enduring ipsilesional evoked activity is more likely to result from the difference in the length of the intervention, the longer of which allows for a reduction in learned disuse of the affected limb (Allred and Jones 2008).

The enduring evoked cortical activity in remote cortical regions that we found elicited by affected limb stimulation at 7weeks post-injury, 4 weeks after intervention, is consistent with an enhanced plastic response, one that has been only found very acutely after injury in this model (Verley et al. 2018). The accompanying FC changes provide ample, evidence for these remote effects, in particular with the reduction in cortico- thalamic and increase in cortico-striatal FC after CIMT compared to noCIMT. Corticothalamic hyperconnectivity is a persistent response both in this model (Harris et al. 2016) and after mild TBI both preclinically and clinically (Li et al. 2023; Woodrow et al. 2023). The enduring alteration in corticothalamic FC associated with earlier CIMT intervention suggests that altered sensory input from both the constrained and force-use limb alters these circuits, possibly at the level of the thalamus, and this may drive the remote changes in limb-evoked cortical activity and FC changes. Emblematic of this in particular is the result of persistent secondary cortex activation after CIMT but not noCIMT, because in the presence of cortical lesioning, the direct corticothalamocortical pathway has been shown to be a potent activator of secondary somatosensory cortex (Theyel, Llano, and Sherman 2010).

The aforementioned circuit changes were only associated with a very mild, temporary trending improvement in injury-affected forelimb use, but the degree of FC change and the endurance of evoked activity do reflect the potential of combinatorial rehabilitation to affect a more marked behavioral improvement. The shaping of forelimb use using CIMT and forelimb reach training (Adkins et al. 2015) would necessarily activate these circuits much more and result in even greater corticothalamic driven plasticity, as shown before at the level of the cortex (Combs et al. 2016). It remains to be seen whether a further combination of direct neuromodulation at the level of the thalamus together with CIMT would potentiate activation of these circuits to an even greater extent, as such driving neural plasticity and enduring improvements in limb function.

